# *Wolbachia* modifies thermal preference in *Drosophila melanogaster*

**DOI:** 10.1101/353292

**Authors:** Amy M. Truitt, Martin Kapun, Rupinder Kaur, Wolfgang J. Miller

## Abstract

Environmental variation can have profound and direct effects on fitness, fecundity, and host symbiont interactions. Replication rates of microbes within arthropod hosts, for example, are correlated with incubation temperature but less is known about the influence of host-symbiont dynamics on environmental preference. Hence, we conducted thermal preference (*T*_*p*_) assays and tested if infection status and genetic variation in endosymbiont bacterium *Wolbachia* affected temperature choice of *Drosophila melanogaster*. We demonstrate that isogenic flies infected with *Wolbachia* preferred lower temperatures compared to uninfected *Drosophila*. Moreover, *T*_*p*_ varied with respect to three investigated *Wolbachia* variants (*w*Mel, *w*MelCS and *w*MelPop). While uninfected individuals preferred 24.4°C, we found significant shifts of - 1.2°C in *w*Mel- and −4°C in flies infected either with *w*MelCS or *w*MelPop. We, therefore, postulate that *Wolbachia*-associated *T*_*p*_ variation within a host species might represent a behavioral accommodation to host-symbiont interactions and trigger behavioral self-medication and bacterial titer regulation by the host.

## INTRODUCTION

Environmental variations through intrinsic (e.g. physiology, reproduction, metabolism) and extrinsic (e.g. food sources, predation risk, immunity) factors impose a strong impact on the fitness of all organisms (e.g., Levins 1968; Endler 1977; 1986; Fox *et al.* 2001). Temperature is one of the most important environmental abiotic factors that affect the physiology and life history traits in many organisms (Huey and Berrigan 2001; Hoffmann 2010; Bozinovic *et al.* 2011; Amarasekare and Savage 2012). Ectotherms, such as terrestrial insects, depend on ambient conditions to maintain their body temperature within a thermoregulatory range (Angilletta *et al.* 2004). For example, thermal preference (*T*_*p*_) in *Drosophila melanogaster,* a dipteran model species of world-wide distribution, varies with geography and elevation, and is thus potentially shaped by selection (Martin and Huey 2008; Dillon *et al.* 2009; Garrity *et al.* 2010; Hoffmann & Sgrò 2011; Huey *et al.* 2012; Rajpurohit and Schmidt 2016). In addition, variation in temperature can have fundamental effects on ecological interactions among organisms and their symbiotic microbes. Titers of endosymbiotic *Wolbachia* bacteria are highly temperature-dependent in various arthropod hosts. For example, some *Wolbachia* strains have increased replication rates at warmer temperatures (Clancy and Hoffmann 1998; Hurst *et al.* 2000; Mouton *et al.* 2006; Correa and Ballard 2012; Strunov *et al.* 2013a), while others are highly sensitive to heat stress (van Opijnen and Breeuwer 1999; Wiwatanaratanabutr and Kittayapong 2009).

Endosymbionts of the genus *Wolbachia* are widespread and found in more than 50% of all investigated terrestrial and some aquatic insects (Zug and Hammerstein 2012; Weinert *et al.* 2015; Sazama *et al.* 2017). *Wolbachia* have garnered extensive interest due to reproductive manipulations they can inflict on their hosts, i.e., inducing parthenogenesis, male killing, feminization, and cytoplasmic incompatibility (CI). By acting as reproductive parasites these bacteria boost their own transmission (reviewed by Werren *et al.* 2008). However, *Wolbachia* can also behave as facultative or obligate mutualists (reviewed by Zug and Hammerstein 2015) by enhancing host fecundity and fitness (Dedeine *et al.* 2001; Hosokawa *et al.* 2010; Miller *et al.* 2010) and by providing protection against RNA viruses (Hedges *et al.* 2008; Teixeira *et al.* 2008; Moreira *et al.* 2009; Osborne *et al.* 2009). Several closely related genetic variants of *Wolbachia* have been isolated from natural and laboratory populations of *D. melanogaster*. *w*Mel, *w*MelCS, and *w*MelPop, which represent three of the most well-studied *Wolbachia* variants in *D. melanogaster* (Riegler *et al.* 2005), cause very weak, if any, CI in their native host (Hoffmann 1988; Reynolds *et al.* 2003; Veneti *et al.* 2003; Fry *et al.* 2004; Yamada *et al.* 2007), but provide virus protection to varying degrees (Chrostek *et al.* 2013; Martinez *et al.* 2014). Both *w*Mel and *w*MelCS infect natural populations of *D. melanogaster.* Historically, *w*MelCS existed globally at higher prevalence, but in the recent past *w*Mel has almost completely replaced the more ancestral *w*MelCS strain in world-wide populations (Riegler *et al.* 2005; Nunes *et al.* 2008; Richardson *et al.* 2012; Ilinsky 2013; Early and Clark 2014). In contrast, *w*MelPop was isolated from a laboratory stock of *D. melanogaster* during a survey of genetic mutations and represents a pathogenic variant of *w*MelCS (Min and Benzer 1997; Richardson *et al.* 2012; Chrostek *et al.* 2013). Depending on rearing temperature, *w*MelPop infections can lead to a strong reduction of host lifespan with respect to uninfected controls (Min and Benzer 1997; McGraw *et al.* 2002; Reynolds *et al.* 2003; Chrostek *et al.* 2013). This detrimental effect is caused by over-
proliferation in host tissues, such as the brain, retina, and muscles (Min and Benzer 1997; Strunov *et al.* 2013b). Importantly, not only *w*MelPop but also its natural predecessor *w*MelCS have significantly higher cellular densities and growth rates than *w*Mel when assayed in the same fly genetic background at 25°C (**Table 1**; Chrostek *et al.* 2013). While high *Wolbachia* densities result in augmented antiviral protection, they also have negative effects by reducing their host’s lifespan. Accordingly, it has been proposed that the higher titer - and hence more costly - *w*MelCS variant was replaced by the low-titer *w*Mel variant in natural *D. melanogaster* populations (Chrostek *et al.* 2013). Thereby, flies infected with the more recent *w*Mel variant have higher fitness due to lower *Wolbachia* titers compared to flies infected with *w*MelCS. Alternatively, the highly protective *w*MelCS variant may have been replaced by *w*Mel independent of the symbiont´s capacity for virus resistance but because of better adaptation to viruses at the host level (Martins *et al.* 2014). In line with this hypothesis, a recent study failed to find correlations between RNA virus prevalence and *Wolbachia* frequency in natural populations of *D. melanogaster* (Webster *et al.* 2015). However, the main causalities explaining the well-documented global almost complete replacement of *w*MelCS by *w*Mel in worldwide populations of *D. melanogaster* remains elusive.

**Table 1.** Comparison of strain type titer levels, growth rates, and effects on host’s lifespan at 25°C.

| **Strain type** | **Relative amount of *Wolbachia*** | **Effects on host's lifespan** |
| --- | --- | --- |
| *w*Mel | Lowest titer level and growth rate | No reduction |
| *w*MelCS | Approximately double the titer level compared to *w*Mel and higher growth rate | Some reduction |
| *w*MelPop | Titer level 20 times higher compared to *w*MelCS | Reduction by approximately half |

Note: Information on titer levels, growth rate, host's lifespan effects for *w*Mel and *w*MelCS from Chrostek *et al.* 2013, information on *w*MelPop's effects on host's lifespan from Reynolds *et al.* 2003.

Host-symbiont conflicts may arise from disparities between physiological requirements of *Wolbachia* and those of their hosts. For example, some insects induce behavioral fever (Louis *et al.* 1986) or behavioral chill (Fedorka *et al.* 2016) as an immune strategy to fight bacterial pathogen infections. Conversely, some bacterial symbionts are known to alter their host’s thermal tolerance range in an adaptive manner (Russell and Moran 2006; Dunbar *et al.* 2007; reviewed by Wernegreen 2012). We, therefore, speculate that additional ecological and behavioral factors, such as host temperature preference, may play a pivotal role in determining *Wolbachia* prevalence and the dynamics of their strain replacement in natural *D. melanogaster* populations.

To test our hypothesis, we conducted laboratory-based temperature preference assays using isogenic *D. melanogaster w*^*1118*^ strains that are either uninfected (*w-*) or infected with one of the three common *Wolbachia* strains *w*Mel, *w*MelCS_b, and *w*MelPop (Teixeira *et al.* 2008; Chrostek *et al.* 2013) and determined if *Wolbachia* affects the temperature preference of its native host *D. melanogaster*. To this end, we built a custom thermal gradient apparatus and determined the temperature preference of replicated fly populations with varying *Wolbachia* infection statuses along the thermal gradient ranging from 17°C to 32°C. Our experiments demonstrate that the temperature preference of *D. melanogaster* is neither sex-nor age-dependent, but is highly dependent on the *Wolbachia* infection status and on the symbiont genotype. Our results provide compelling evidence that *Wolbachia* infections can affect host thermal preference behavior, at least under strict laboratory conditions in *D. melanogaster* strains.

## RESULTS

To determine whether *T*_*p*_ of adult *D. melanogaster* varies with *Wolbachia* infection status and *Wolbachia* genotype, we conducted lab-based experiments using a custom-built temperature gradient apparatus for assaying flies of the isogenic lab-strain *w*^*1118*^ that were either uninfected (*w*-) or infected with one of the *Wolbachia* strains *w*Mel, *w*MelCS, or *w*MelPop (**Supporting Information Fig. S1-4**). We first investigated whether age (3-4, 5-7 or 10-14 days post eclosion) and *Wolbachia* infections, or sex (males or females) and *Wolbachia* infections had an influence on *T*_*p*_ by means of two-way mixed-effect Poisson regressions. We neither found significant effects of age or sex nor significant interactions of either factor with *Wolbachia* infections (see **Fig. 1A+B** and **Table 2A+B** and **Supporting Information Fig. S5**; Poisson regression: *P* > 0.05 for factors age and sex and both interaction terms, respectively). In contrast, both two-way regressions revealed highly significant effects of *Wolbachia* infections on *T*_*p*_ (Poisson regression P < 0.001 for factor *Wolbachia* in both analyses). Since both aforementioned analyses were carried out on different subsets of the data which did not include all four infection types (*w*-, *w*Mel, *w*MelCS and *w*MelPop), we further investigated all data jointly irrespective of sex and host age and evaluated the effect of symbiont genetic variation on *T*_*p*_ by means of post-hoc pairwise comparisons based on Tukey’s honestly significant differences (HSD). We found that temperature preference of *D. melanogaster* strongly depended on (1) the infection status of the flies and (2) on the *Wolbachia* strain used for infections: Uninfected flies (*w*-) exhibited the highest mean *T*_*p*_ at 24.4°C (Median: 25°C; Mode: 26°C), while *w*Mel-infected flies preferred average temperatures at 23.2°C (Median: 24°C; Mode: 24°C), which is 1.2°C lower than uninfected (*w*-) flies. In contrast, flies infected with *w*MelCS or *w*MelPop showed highly similar thermal preferences at 20.6°C and 20.5°C (Median: 19°C and Mode: 18°C for both) respectively, which were both approximately 4°C lower than to *w*-(see **Fig. 1C**, **Table 2C** and **Table 3**).

**Figure 1:**
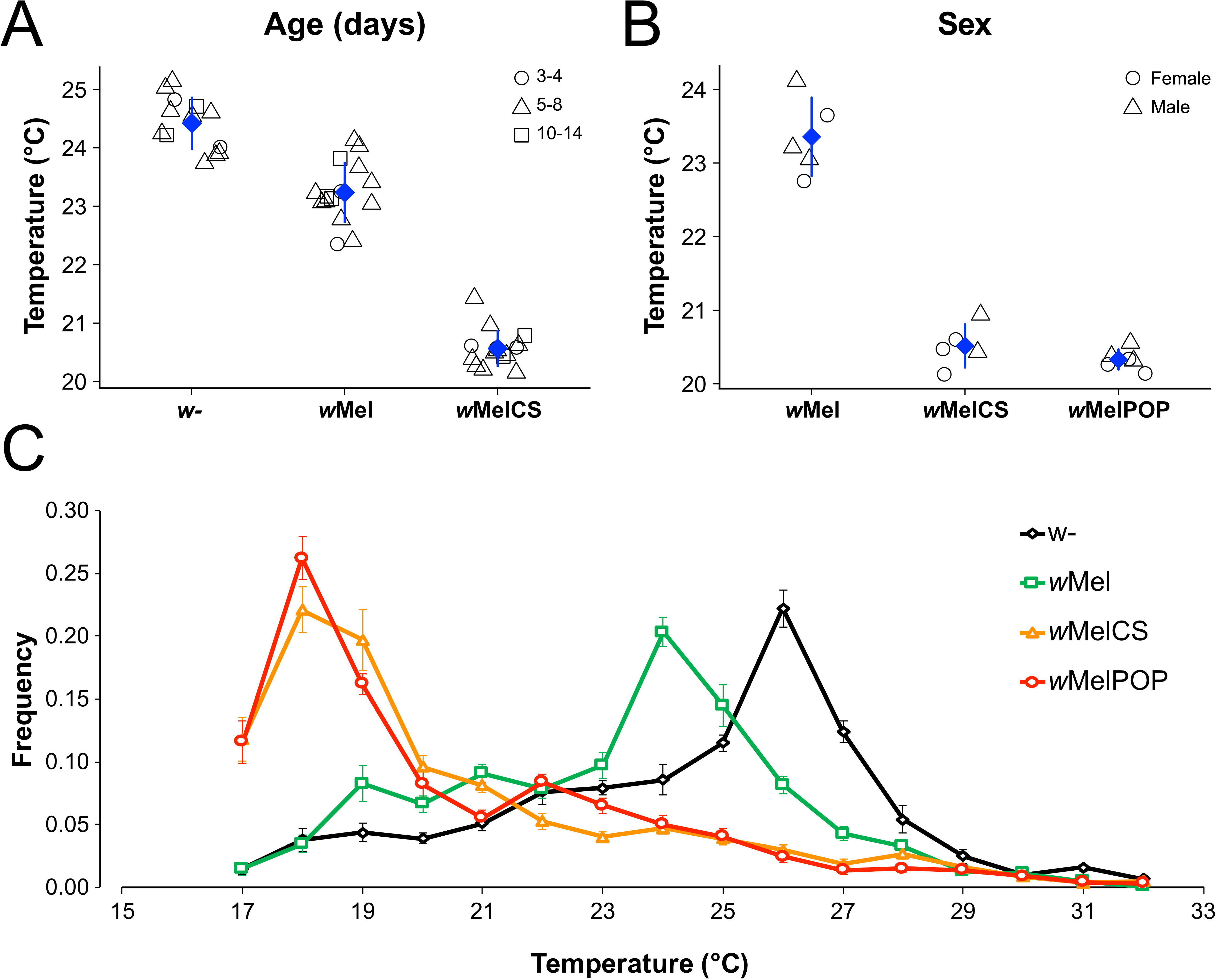
Thermal preference of *Drosophila* with and without *Wolbachia* infections. Panels A and B show average *T*_*p*_ (blue diamonds) with respect to age (3-4, 5-7 or 10-14 days post eclosion, n=4370 excluding flies infected with *w*MelPop) and sex (male or female; n=1718, excluding uninfected flies), respectively. Each symbol represents the average *T*_*p*_ for a replicate at a given factor level of either age (circle: 3-4 days, triangle: 5-8 days and square: 10-14 days) or sex (circle: females, triangle: males). Panel C shows line plots with relative proportions of flies observed at a given temperature. Each line represents the average proportion of flies which were either uninfected (*w*-; black), or infected with *w*Mel (red), *w*MelCS (blue) or *w*MelPop (green). The error bars represent standard errors for average frequencies at a given temperature across all replicated experiments carried out for each infection type. We found that infected flies exhibit significantly lower thermal preference compared to uninfected flies.

**Table 2.** Table showing the results of three analyses based on generalized linear mixed models with a Poisson error structure to account for the statistical properties of count data. The columns show ID’s for the different analyses (A-C), the models, the individual factors and interactions tested, the samples size, the degrees of freedom for the *χ*^2^ test of the analysis of deviance, the *χ*^2^ value and the corresponding *P*-value. Note that analyses with significant effects after Bonferroni correction (adjusted *α* = 0.017) are highlighted in bold.

| **Analysis** | **Model** | **Factor** | ***n*** | **Df** | **$\chi^2$** | ***P*-value** |
| --- | --- | --- | --- | --- | --- | --- |
| **A** | ***wol + age + wol x age*** | ***wol*** | **4370** | **6** | **119.71** | **1.87E-23** |
| A | *wol + age + wol x age* | *age* | 4370 | 6 | 2.10 | 0.91 |
| A | *wol + age + wol x age* | *wol x age* | 4370 | 4 | 1.31 | 0.86 |
| **B** | ***wol + sex + wol x sex*** | ***wol*** | **1718** | **6** | **61.19** | **2.58E-11** |
| B | *wol + sex + wol x sex* | *sex* | 1718 | 4 | 2.12 | 0.71 |
| B | *wol + sex + wol x sex* | *wol x sex* | 1718 | 3 | 1.90 | 0.59 |
| **C** | ***wol*** | ***wol*** | **5717** | **3** | **168.69** | **2.44E-36** |

**Table 3.** Table showing z-values from post-hoc pairwise comparisons with Tukey’s HSD for the factor Wolbachia (Analysis C; see Experimental Procedures) with four levels (non-infected, wMel, wMelCS, and wMelPop). Bold type indicates significance after Bonferroni correction (adjusted *α*’= 0.017). * p < 0.05; ** p < 0.01; *** p < 0.001.

|  | *w*- | *w*Mel | *w*MelCS | *w*MelPop |
| --- | --- | --- | --- | --- |
| *w*- | - |  |  |  |
| *w*Mel | -6.76*** | - |  |  |
| *w*MelCS | -21.93*** | -15.49*** | - |  |
| *w*MelPop | -21.5*** | -15.35*** | -0.6 | - |

## DISCUSSION

In this study, we, for the first time, investigated the relationship between temperature preference of *D. melanogaster* and *Wolbachia* infection under laboratory conditions. Using a custom-built thermal gradient apparatus, we conducted temperature preference assays and showed that the *T*_*p*_ of *D. melanogaster* is shifted to lower temperatures when flies are infected with *Wolbachia*. Uninfected *D. melanogaster* flies preferred an average temperature of 24.4°C, whereas *w*Mel-infected flies preferred 23.2°C, and both *w*MelCS-and *w*MelPop-infected flies preferred 20.6°C and 20.5°C respectively.

*T*_*p*_ can vary significantly between populations of the same species (Matute *et al.* 2009; Rajpurohit and Schmidt 2016) and can have profound effects on immune function, fitness, and fecundity (Huey and Berrigan 2001; Martin and Huey 2008; Hoffmann 2010). Recent population analyses of *Wolbachia* and mitochondria from *D. melanogaster* have provided evidence that over the past few thousand years, the *w*MelCS variant is being globally replaced by the *w*Mel-variant (Riegler *et al.* 2005; Nunes *et al.* 2008; Richardson *et al.* 2012; Early and Clark 2013). Rare cases of the *w*MelCS infection type were recently detected in the wild (Nunes *et al.* 2008; Ilinsky 2013), thus replacement by *w*Mel is still incomplete. Although the reason for the worldwide turn-over remains elusive, it has been hypothesized that *w*Mel, which persists in hosts at significantly lower densities than *w*MelCS at 25°C (Chrostek *et al.* 2013), has better adapted to *D. melanogaster.* Accordingly, *w*Mel infections are less costly to the host compared to the more ancestral *w*MelCS variant (Chrostek et al. 2013; reviewed by Miller 2013).

Insects can actively reduce or avoid costs of potentially fitness-reducing symbionts or parasites by behavioral adjustments such as changing egg deposition (Kacsoh *et al.* 2013) or mating behavior (reviewed by Wedell 2013). We find compelling evidence for *Wolbachia*-induced behavioral changes in host *T*_*p*_, which may provide an alternative explanation for the recent global replacement of *w*MelCS by *w*Mel independent of density costs or anti-viral effects: we propose that *w*Mel is less costly for the host than *w*MelCS-infections because flies harboring *w*Mel exhibit thermal preferences that are closer to uninfected flies under natural conditions compared to flies infected with *w*MelCS. Since *Drosophila* development is strictly temperature dependent (approximately 14 days of egg-to-adult development at 20°C and 9 days at 24°C; Ashburner 1989), flies infected with *w*Mel should have shorter generation times and thereby produce more generations per year resulting in higher net fecundity compared to *w*MelCS infected flies.

Small fluctuations in temperature can cause considerable modifications to host-symbiont interactions (Blanford and Thomas 1999). Pathogenicity of *w*MelPop is attributed to its active proliferation in host tissues at temperatures ≥ 19°C. The increase of *w*MelPop density confers strong anti-viral protection but leads to a significant reduction in host lifespan at 25°C (Chrostek *et al.* 2013). However, at temperatures < 19°C, pathogenicity of *w*MelPop is eliminated (Reynolds *et al.* 2003). Similarly, but less dramatically *w*MelCS, the progenitor of *w*MelPop, is also costly by reducing host lifespan due to high symbiont densities at 25°C (Chrostek *et al.* 2013). We, therefore, speculate that the adjustment of lower temperature preference in *D. melanogaster* as a response to the *w*MelCS and *w*MelPop infections represents a physiological self-medicating behavior or behavioral chill (Fedorka *et al.* 2016) to attenuate the fitness costs associated with deleterious effects of *Wolbachia* over-proliferation and high cell densities (Chrostek *et al.* 2013; Strunov *et al.* 2013a; Strunov *et al.* 2013b).

*Wolbachia’*s ability to provide anti-viral protection to their hosts has emerged as the most promising approach to combatting insect-vector borne pathogens that pose serious health risks to humans, such as dengue fever and Zika (Moreira *et al.* 2009; Iturbe-Ormaetxe *et al.* 2011; Dutra *et al.* 2016). However, because the strength of anti-viral protection is associated with higher *Wolbachia* densities (Chrostek *et al.* 2013; Martinez *et al.* 2014) and bacterial titers are a temperature sensitive trait (Hoffmann *et al.* 1990; Reynolds *et al.* 2003; Mouton *et al.* 2006; Mouton *et al.* 2007; Bordenstein & Bordenstein 2011; Correa and Ballard 2012; Chrostek *et al.* 2013; Strunov *et al.* 2013a; Strunov *et al.* 2013b; Murdock *et al.* 2014; Versace *et al.* 2014), it is feasible that under certain thermal conditions such as lower environmental temperatures, *Wolbachia-*induced virus protection could be attenuated or absent (Chrostek 2014). Furthermore, our findings, as demonstrated in a highly inbred lab strain of *D. melanogaster,* need to be tested first in different host backgrounds, which are naturally or artificially infected with the endosymbiont.

In conclusion, we present experimental support for a potential ecological conflict between host and symbiont that may have profound effects on host physiology. Our results provide a novel conceptual platform from which to further investigate host temperature preference, or behavioral chill, in other *Wolbachia*-infected insect hosts. Future studies should examine if host temperature preference has a direct impact on *Wolbachia* density regulation. Additionally, it is important to determine any effects that host *T*_*p*_ has on the strength of anti-viral protection that *Wolbachia* provide to some hosts.

## EXPERIMENTAL PROCEDURES

### Fly Lines

For all assays, we used *D. melanogaster* without *Wolbachia* (*w*-) as well as flies infected with one of three genetic variants of the *Wolbachia w*Mel-strain; *w*Mel, *w*MelCS_b, and *w*MelPop all set in the DrosDel *w*^*1118*^ isogenic background, which were kindly provided by Luis Teixeira and previously described by Teixeira *et al.* (2008) and Chrostek *et al.* (2013).

We used biological replicates of approximately 30 flies per vial, independently rearing each vial of flies at 25°C, in a 12:12 light - dark cycle with constant 45% humidity. Flies were raised on *Drosophila* Formula 4-24^®^ Instant Medium (Carolina^®^, NC) that was supplemented with fresh yeast. Approximately equal numbers of male and female flies were used in each assay except for assays that explicitly tested sex-class *T*_*p*_ differences (see **Supporting Information Table S1 and Supporting Information File 1**). In addition to testing for sex-class *T*_*p*_ differences, we performed assays to test for age-specific *T*_*p*_ differences, thus all fly lines were segregated into three age-classes – 3-4 days, 5-7 days, and 10-14 days post eclosion. Due to fitness costs to the host associated with infection by *w*MelPop at 25°C, possibly due to the onset of the life reducing phenotype (Min and Benzer 1997) or increase in copy numbers of the Octomom repeat (Chrostek and Teixeira 2015), our *w*MelPop-infected fly line did not produce enough flies to conduct all three age-class assays. Therefore, we excluded *w*MelPop from the statistical analyses of age-specific effects (see **Supporting Information Table S1** and the description of statistical analyses).

### Genotyping of *Wolbachia* strains

Genome sections that contain hypervariable loci or hypervariable regions covering tandem repeats were used as genetic markers to differentiate *Wolbachia* strains and strain variants (O’Neill *et al.* 1992; Werren *et al.* 1995; Zhou *et al.* 1998; Riegler *et al.* 2012). To confirm *Wolbachia-*infection status, we performed diagnostic PCR amplification using primers for a gene that encodes the *Wolbachia* surface protein, *wsp* (Jeyaprakash and Hoy 2000), and for an intergenic region with 141bp tandem repeats, VNTR-141 loci (Riegler *et al.* 2005). The PCR reactions for *wsp* amplification were carried out in a total volume of 10μl containing 2μl Promega 5x Green GoTaq buffer, 4mM Promega MgCl_2_, 0.8μM of forward and reverse primers, 35μM of each dNTP, 0.04 U Promega GoTaq DNA Polymerase, and 1μl of genomic DNA template. Diagnostic VNTR-141 PCR reactions were each a total of 10μl comprised of the following: 2μl Promega 5x Green GoTaq buffer, 1.5mM Promega MgCl_2_, 0.3μM of forward and reverse primers, 35μM of each dNTP, 0.04 U Promega GoTaq DNA Polymerase, and 1μl of genomic DNA template. PCR products were visualized on a 1% agarose gel. Presence/absence of the *wsp* signal and the size of the diagnostic VNTR-141 locus confirmed their respective infection type (Riegler *et al.* 2012). The proper infection status of the *w*MelPop isoline was verified by assaying flies for early mortality at 29°C.

### Thermal gradient apparatus

Temperature preference assays were performed using a custom made thermal gradient apparatus that allowed the flies to move in a three-dimensional space (adapted from Rajpurohit and Schmidt 2016; **Supporting Information Fig. S2**). An aluminum rod (length 74.93cm, diameter 3.02cm; Part #R31-316 Metals Depot, Winchester, KY) was encased within a 58.76cm long and 6.35cm inside diameter polycarbonate tube, creating an enclosed chamber allowing for three-dimensional movement. Constant voltage was applied to Peltier devices on each end of the aluminum rod to create a temperature gradient inside the thermal preference chamber. Temperatures along the gradient were measured at seven points that were 8.39cm apart using K-type thermocouples and two four-channel thermocouple recorders. We recorded temperatures on the aluminum rod and inside polycarbonate tube surfaces (bottom, top, and mid-point between the top and bottom surfaces; **Supporting Information Fig. S3**). The average temperatures from each thermocouple point on all surfaces from 57 different assays are depicted in **Supporting Information Fig. S1**. Mean temperatures increased linearly and ranged from 12°C at the coldest point to 40°C at the hottest point of the aluminum rod, 58.76 cm distance (**Supporting Information Fig. S4**). Along the aluminum rod, for every 4.2cm from cold to hot, the temperature increased by 2°C. Temperatures along each of the measured polycarbonate tube surfaces (bottom, mid-point, and top) increased 1°C every 4.2cm from cold to hot. The gradient reached thermal stability after approximately 20 minutes and remained stable for at least 3 hours. Assays were conducted once the device had attained thermal stability.

### Thermal preference assays

All assays were conducted in a room with a constant temperature of 24°C and constant 40% humidity. During several trial runs, we established that 75-100 flies for each assay resulted in distributions along the thermal gradient that avoided over-crowding in preferred temperature ranges, eliminating potential counting errors during analysis. Flies were introduced by aspiration into the thermal gradient chamber through a small hole located halfway along the top of the polycarbonate tube, where the temperature consistently averaged 25°C. Flies used for thermal preference assays were never anesthetized because of the strong effects from CO_2_ treatment on *Drosophila* behavior (Barron 2000). Each assay was conducted for thirty minutes. Between assays, the temperature gradient chamber was taken apart and thoroughly cleaned to avoid contamination from any pheromone particles. All aluminum parts were cleaned using 95% ethanol. Because ethanol and polycarbonate are chemically incompatible, the polycarbonate tube and end caps were cleaned using hot water and soap, followed by a four-minute rinse with hot water to ensure that surfaces were free of soap residue.

### Data collection

Using three GoPro HERO3+ cameras, we collected data for each assay in the form of digital images. To capture images of the entire thermal gradient and the flies within it, we mounted the cameras above, lateral to, and below the apparatus, capturing images every 30 seconds for the duration of each treatment (30 minutes). Images were analyzed using Adobe Photoshop CS6. All 60 images from each assay were reviewed, from which we determined that A) the flies were highly active, retaining the ability to relocate as necessary, for the entire assay, and B) after being introduced to the thermal gradient, actively flew around for up to 15 mins before they settled on either the aluminum rod or polycarbonate tube surfaces. Therefore, we selected images for analysis of fly distribution at the 20-minute time point as representative of the 30-minute experiment. For each assay, we manually counted flies and marked the location of flies on a custom grid that delineated gradient surfaces and surface temperatures.

### Statistical analyses

We calculated generalized linear mixed models (GLMM) with a Poisson error structure using the *R* (R Development Core Team 2009) package *lme4* (Bates *et al.* 2015) to account for the statistical properties of count data from flies observed at different temperatures. To test for significance of a given predictor variable, we compared the full model including all factors to a reduced model excluding the given factor by analysis of deviance with *χ*^2^ tests using the *R* function *anova* (see **Supporting Information File 1** for full *R* code).

At first, we excluded flies infected with *w*MelPop, since we failed to obtain sufficient flies to test for age-specific *T*_*p*_ at all three age-classes (3-4 days, 5-7 days and 10-14 days post eclosion; **Supporting Information Table S1**) and tested for age-and *Wolbachia*-specific differences in thermal preference with a two-way GLMM of the form: *T*_i_ = *wol* + *age* + *wol* × *age* + *Rep* + *ɛ*_i_. Here, *T* is the continuous response variable “Temperature”, *age* is a nominal fixed factor with three levels each (*age*: 3-4 days, 5-7 days and 10-14 days post eclosion), *wol* is a nominal fixed factor “*Wolbachia*” with three levels (un-infected, *w*Mel and *w*MelCS), *wol* × *age* is the interaction term, *Rep* is a nominal random factor “Replicate” for replicate trials and *ɛ*_i_ is the error (**Table 2A**, **Fig. 1A**). In a complementary analysis, we removed all flies of the age class 3-4 days and repeated the abovementioned analysis including all *Wolbachia* strains on two age classes (5-7 days and 10-14 days post eclosion) only. This latter analysis yielded qualitatively similar results to the former analysis including all age classes without *w*MelPop (**Supporting Information Table S2**).

Next, we censored flies with undetermined sex status and excluded uninfected flies (*w*-), since we failed to obtain sufficient replication to test for male-specific *T*_*p*_ for uninfected flies (**Supporting Information Table S1**). We then tested for sex- and *Wolbachia*-specific differences in thermal preference with a two-way GLMM of the form: *T*_i_ = *wol* + *sex* + *wol* × *sex* + *Rep* + *ɛ*_I_ Here, *T* is the continuous response variable “Temperature”, *sex* is a nominal fixed factor with two levels (male and female), *wol* is a nominal fixed factor “*Wolbachia*” with three levels (*w*Mel, *w*MelCS, and *w*MelPop), *wol* × *age* is the interaction term, *Rep* is a nominal random factor “Replicate” for replicate trials and EI is the error (**Table 2B; Fig. 1B**).

Finally, we included all flies, irrespective of age and sex status, and tested for the effect of infection status and *Wolbachia* strain variation on thermal preference with a GLMM of the form: *T*_i_ = *wol* + *Rep* + *ɛ*_i_, where *T* is the continuous response variable “Temperature”, *wol* is a nominal fixed factor “*Wolbachia*” with four levels (un-infected, *w*Mel, *w*MelCS, and *w*MelPop), *Rep* is the nominal random factor “Replicate” and E_i_ is the error (**Table 2C; Fig. 1C**). Here, we further tested for significant pair-wise comparisons among the level of the factor “*Wolbachia*” with Tukey’s honestly significant difference (HSD) post-hoc tests using the *R* package *multcomp* (**Table 3**). We conservatively applied Bonferroni corrections to the *α* threshold (*α*’= 0.05/3 = 0.017) to account for multiple testing.

## Acknowledgement

The authors want to thank Luis Teixeira for providing the *Drosophila melanogaster* white-isogenic DrosDel *w*^*1118*^ lines infected with *w*Mel, *w*MelCS_b and *w*MelPop. The authors declare no conflict of interest.

## Author contributions

A.T. and W.J.M. conceived and planned the study, A.T. performed the experiment, A.T., R.K., and M.K. analyzed the data and A.T., W.J.M., and M.K. wrote the paper.

